# Inhibition of farnesyltransferase activity diminishes hematopoietic stem cell *ex vivo* expansion ability

**DOI:** 10.1101/2025.10.30.685582

**Authors:** Carli Newman, Isabella M. Alves, Matthew W. Hagen, Rachel Wellington, Brandon Hadland, Anita M. Quintana, Christina Marie Termini

**Affiliations:** Translational Science and Therapeutics Division, Fred Hutchinson Cancer Center, Seattle, WA, USA; Human Biology Division, Fred Hutchinson Cancer Center, Seattle, WA, USA; Institute of Stem Cell and Regenerative Medicine, University of Washington, Seattle, WA, USA; Division of Hematology Oncology and Bone Marrow Transplant, Department of Pediatrics, University of Washington, Seattle, WA, USA; Department of Biology, University of Texas at Arlington, Arlington, TX, USA; Department of Laboratory Medicine and Pathology, University of Washington, WA, USA

## Abstract

The rationale underlying this work is that elucidating the contribution of farnesyltransferase activity to hematopoietic stem cell expansion *ex vivo* will provide knowledge needed to better support hematopoietic stem cell expansion techniques for clinical applications that rely on this approach, like hematopoietic cell transplants and gene therapy. We discovered that pharmacological inhibition of farnesyltransferase activity with lonafarnib substantially diminished the *ex* vivo expansion potential of human and mouse hematopoietic stem cells, highlighting that hematopoietic stem cells rely on isoprenoids for their *ex vivo* maintenance.

## MAIN TEXT

Due to their robust regenerative properties, hematopoietic stem cells (HSCs) are used in cell and gene therapy approaches to boost or restore the function of the hematopoietic system in patients [1]. For example, hematopoietic cell transplants are used to treat patients with blood cancers and immune disorders [2]. More recently, HSCs have been leveraged for gene-editing approaches to correct defective hematopoietic cells [3]. While the potential therapeutic applications of HSCs are vast and growing, a barrier to obtaining sufficient HSC numbers to make these technologies accessible to all still exists. HSC expansion *ex vivo* can be used to increase the number of hematopoietic cells generated from HSCs to procure sufficient cell numbers for cell and gene therapy approaches [4, 5]. Several cell-intrinsic pathways that support HSC expansion have been identified, such as those enacted by stem cell factor (SCF), thrombopoietin (TPO), and the FMS-like tyrosine kinase 3 ligand (FLT3L) [6]. Compared to proteins, our understanding of the roles of lipids in regulating HSC *ex vivo* expansion ability is much more limited [7].

The mevalonate pathway (MVA) generates lipids through enzymatically controlled reactions [8] (**Figure 1A**). Farnesyl-pyrophosphate is a metabolic intermediate that can be further converted to generate either cholesterol or isoprenoids, like geranylgeranyl pyrophosphate. Cholesterol supports plasma membrane structure and function and is a precursor to key biological components, like hormones. Dysregulated cholesterol homeostasis disrupts hematopoietic stem/progenitor cell mobilization and repopulation potential, highlighting the importance of lipids in hematopoietic cell functions [9, 10]. Isoprenoids serve as substrates for proteins to be post-translationally modified via farnesylation, a process mediated by farnesyltransferases. Farnesylation of molecules paramount to HSC functions, like Ras and Rho GTPases, is known to drive their activity [11, 12]. Further, pharmacological inhibition of farnesylation caused developmental anemia in zebrafish [13]. Given that HSCs depend on proteins that are known to be farnesylated [14, 15], we hypothesized that inhibiting protein farnesylation would impair HSC functions. Here, we determined that inhibition farnesylatransferase activity reduces HSC *ex vivo* expansion ability and promotes cell death.

**Figure 1:**
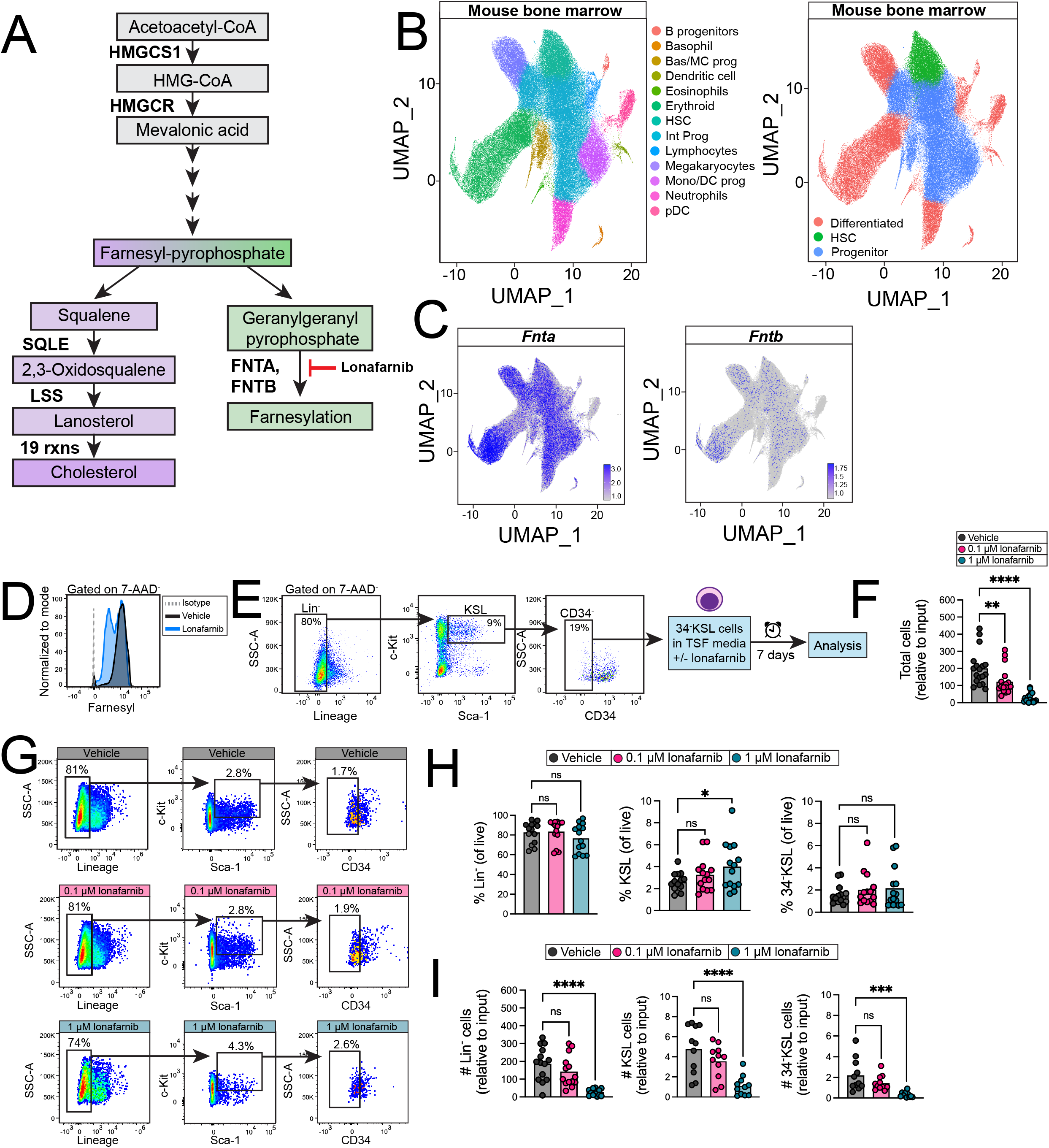
Farnesyltransferase inhibition reduces mouse HSC expansion *ex vivo*. (A) Diagram depicting the major steps in the mevalonate pathway. **(B)** Uniform Manifold Approximation and Projection (UMAP) visualization of bone marrow niche cells from dataset analyses in PMID: 38183977 [16]. Each color represents a different annotated cell population and **(C)** relative expression of *Fnta* and *Fntb* with 10-90^th^ percentile of expression. **(D)** Histogram depicting farnesyl expression in Lin-cells treated with lonafarnib of vehicle for 24 hours. **(E)** Gating strategy used to isolate 34^-^KSL cells from the bone marrow of adult C57BL/6 mice and culturing strategy; frequencies on plots are relative to parent population. **(F)** Total cells were quantified at day +7 after culturing 34^-^KSL cells with lonafarnib in TSF media and normalized to input cell number. **(G)** Representative gating strategy used to analyze hematopoietic stem/progenitor cell populations. Frequencies show frequencies relative to total cells in the well. **(H)** HSPC frequencies at day +7 relative to live cells. **(I)** The numbers of HSPCs per well at day +7 after lonafarnib treatment, shown normalized to input. (*n=3-5 biological replicates per experiment from n=4 independent experiments experiment, error bars show SEM, statistics show one-way ANOVA followed by Holm-Sidak’s multiple comparisons tests*).

We first analyzed the expression of transcripts encoding for genes that catalyze farnesylation in mouse bone marrow (BM) hematopoietic cells (**Figure 1B**) using a published dataset [16]. *Fnta* (farnesyltransferase/geranylgeranyltransferase type-1 subunit alpha) and *Fntb* (farnesyltransferase subunit beta) were expressed broadly in the BM (**Figure 1C**). These data indicate that mouse BM hematopoietic cells genes that encode for farnesyltransferases, but their expression is not restricted to HSCs or a particular hematopoietic lineage.

Since lipids have been shown to regulate HSCs [9, 10] and genes encoding farnesyltrasnferases are broadly expressed in HSCs and the mouse BM (**Figure 1A-C**), we analyzed the function of farnesyltransferase activity in HSC expansion *ex vivo*. For these studies, we used lonafarnib, a pharmacological inhibitor of farnesyltransferase activity, which we validated to deplete farnesyl levels in hematopoietic cells (**Figure 1D**). Using fluorescence-activated cell sorting we isolated BM Lineage^-^c-Kit^+^Sca-1^+^ (KSL) CD34^-^ HSCs (herein referred to as 34-KSL HSCs) from adult wild-type C57BL/6 mice. Isolated 34-KSL cells (**Figure 1E**) were cultured cells in cytokine-enriched expansion media containing TPO, SCF, and FLT3L (TSF media), supplemented with vehicle or increasing lonafarnib concentrations.

Lonafarnib significantly reduced HSC expansion in a concentration dependent manner compared to vehicle (**Figure 1F**). Lineage negative (Lin^-^) HSPC and 34^-^KSL HSC frequencies were unchanged (**Figure 1G-H**), while KSL cell frequency was increased (**Figure 1H** middle panel) upon lonafarnib treatment. These data suggest that lonafarnib treatment has a minor impact on hematopoietic stem and progenitor cell balance upon expansion. However, lonafarnib treatment significantly reduced the absolute numbers of Lin^-^, KSL, and 34^-^ KSL cells produced relative to input compared to vehicle treatment (**Figure 1I**). Further, lonafarnib treatment reduced 34^-^KSL cells relative to input, suggesting an accelerated depletion of the HSC pool compared to vehicle control. These data suggest that inhibition of farnesyltransferase activity reduces hematopoietic expansion ability.

Detailed methods used throughout our study are outlined in the **Supplementary Methods** and resources are detailed in **Table 1**. To determine how our findings align with human systems, we queried published single-cell RNA sequence datasets of human BM hematopoietic cells (**Figure 2A**) [17]. Similar to mouse, human hematopoietic cells also express *FNTA* and *FNTB* broadly (**Figure 2B**). Therefore, genes encoding farnesyltransferases are expressed in human BM hematopoietic cells but not enriched in HSCs.

**Figure 2:**
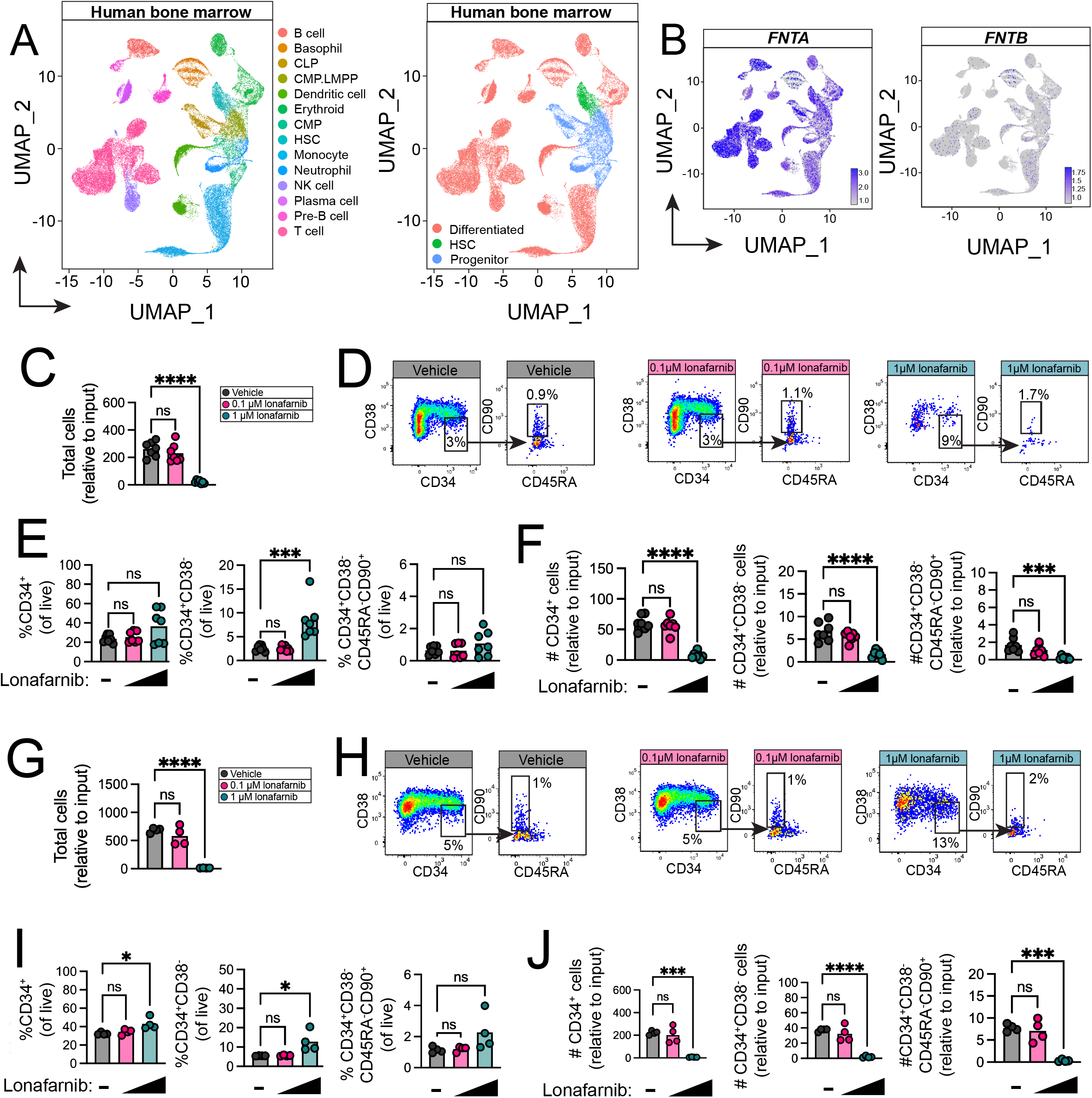
Farnesyltransferase inhibition reduces human HSC expansion *ex vivo*. **(A)** UMAP visualization of human bone marrow cells from dataset analyzed in PMID: 31792411 [17]. **(B)** Relative expression of *FNTA* and *FNTB* in human bone marrow cells with 10-90^th^ percentile of expression. (**C)** Human bone marrow CD34^+^ cells were cultured in TSF media with or without lonafarnib, and total cells were quantified. **(D)** Representative gating strategy used to analyze human HSPCs at day +7 after culturing with lonafarnib. **(E)** The frequency of HSPCs relative to live cells per well at day +7 after culturing in TSF enriched with lonafarnib and **(F)** the number of bone marrow HSPCs per well quantified and normalized to input cell number. **(G)** Human cord blood CD34^+^ cells were cultured in TSF media with or without lonafarnib, and total cells were quantified. **(H)** Representative gating strategy used to analyze human HSPCs at day +7 after culturing with lonafarnib. **(I)** The frequency of HSPCs relative to live cells per well at day +7 after culturing in TSF enriched with lonafarnib and **(J)** the number of cord blood HSPCs per well quantified and normalized to input cell number. (*n=3-5 biological replicates per experiment from n=2 independent experiments with unique donors used in each experiment, error bars show SEM, statistics show one-way ANOVA followed by Holm-Sidak’s multiple comparisons tests*).

Human BM CD34^+^ HSPCs cultured in TSF media enriched with 1µM lonafarnib exhibited a more than tenfold decrease in hematopoietic cells per well overall compared to vehicle-treated cells (**Figure 2C**). Analysis of immunophenotypically-defined HSPCs after lonafarnib treatment revealed a significant increase in the percent of CD34^+^CD38^-^ cells per well compared to vehicle treatment (**Figure 2D-E**). However, lonafarnib treatment reduced the number of CD34^+^, CD34^+^CD38^-^, and CD34^+^CD38^-^CD45RA^-^CD90^+^ cells per well compared to vehicle (**Figure 2F**). We performed analogous studies using human umbilical cord blood (CB) CD34^+^ cells. We detected similar trends as in BM, with more than a tenfold reduction in overall cell expansion at day +7 upon 1 µM lonafarnib compared to 0.1 µM lonafarnib or vehicle control (**Figure 2G**). Lonafarnib treatment increased CD34^+^, CD34^+^CD38^-^, and CD34^+^CD38^-^CD45RA^-^CD90^+^ cell frequencies (**Figure 2H-I**). However, absolute numbers of CD34^+^, CD34^+^CD38^-^ and CD34^+^CD38^-^CD45RA^-^CD90^+^ HSPCs were decreased upon 1 µM lonafarnib treatment (**Figure 2J**). Together, these data show that inhibition of farnesyltransferase activity reduces adult and primitive human HSPC cell expansion potential overall and depletes the HSPC pool upon *ex vivo* culture.

To evaluate the effect of lonafarnib treatment on hematopoietic progenitor cell differentiation and proliferation, we used colony forming unit (CFU) assays. Mouse 34^-^KSL cells were cultured in methylcellulose containing vehicle or lonafarnib, and we quantified hematopoietic colony formation. There was a significant decrease in colony numbers overall upon lonafarnib treatment (**Figure 3A**). This decrease in colony forming potential occurred within the granulocyte-macrophage (GM) progenitor, while CFU granulocytes, erythrocytes, monocytes, and macrophages (GEMM), and CFU erythroid and burst-forming unit erythroid (BFU-E) colonies were unchanged. We similarly quantified the CFU potential of human BM and CB CD34^+^ cells grown in MethoCult supplemented with vehicle or lonafarnib. In human BM CD34^+^ cells there was a significant reduction in CFU-GM and BFU-E colony formation in the presence lonafarnib compared to vehicle (**Figure 3B**). Meanwhile, cord blood CD34^+^ cells treated with lonafarnib exhibited depleted CFU-GM potential compared to vehicle (**Figure 3C**). These data indicate that inhibiting farnesyltransferase activity reduces mouse and human hematopoietic progenitor cell potential *in vitro*. Further, these findings highlight distinct effects within the erythroid progenitor compartment in human bone marrow CD34^+^ cells, which have have depleted erythroid-committed progenitor cell functionality in the presence of lonafarnib. These findings align with prior studies showing that inhibiting farnesyltransferase activity causes anemia in zebrafish and highlight the potential for hematopoietic progenitor cells to contribute to the detected effects [13].

**Figure 3:**
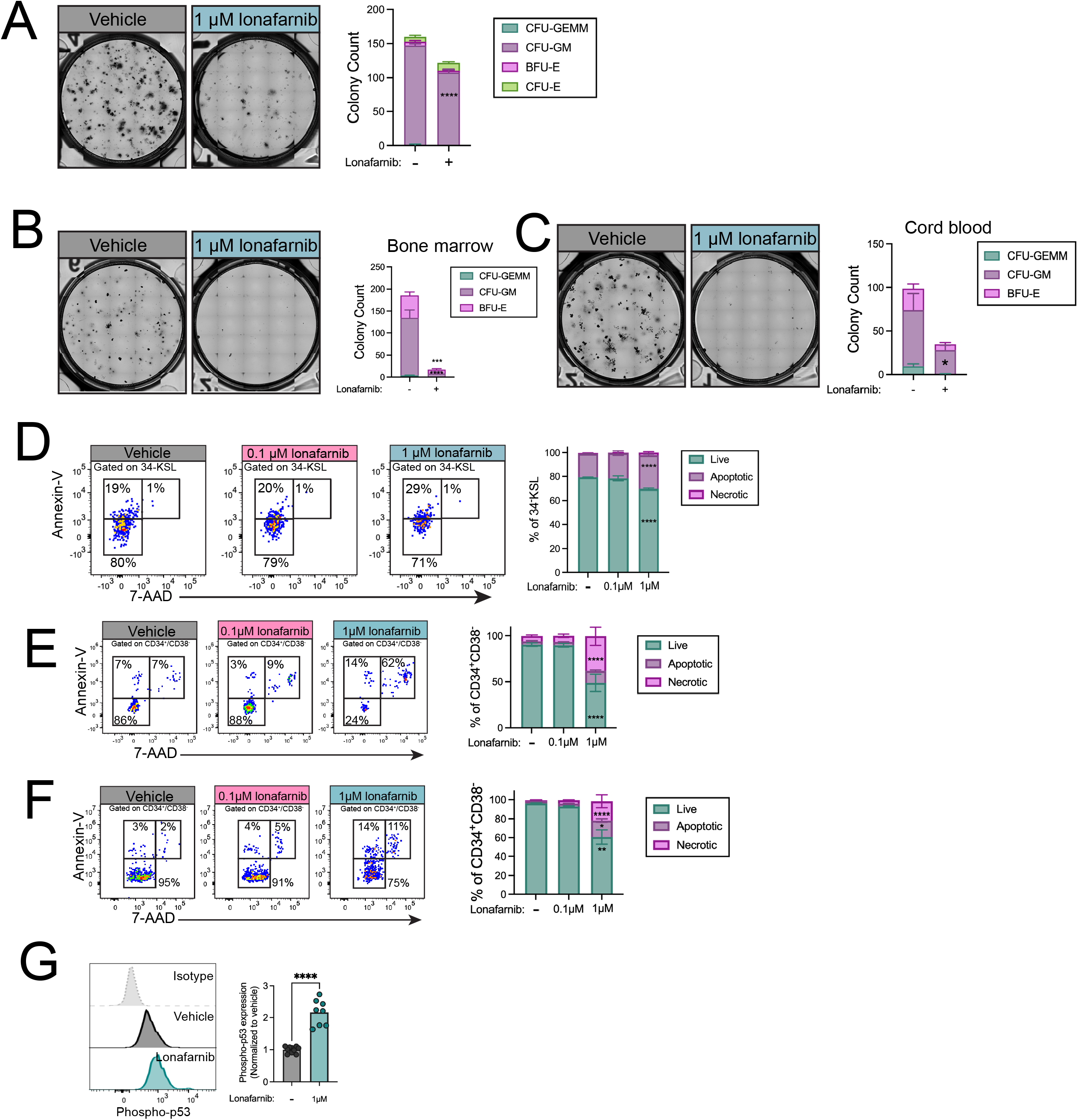
Lonafarnib treatment depletes hematopoietic colony formation and causes cell death. **(A)** STEMVision images showing colony formation and quantification of mouse hematopoietic colonies formed from 34-KSL cells grown in the presence of lonafarnib or vehicle. STEMVision images depicting colony formation from **(B)** bone marrow or **(C)** cord blood CD34+ cells grown with lonafarnib or vehicle. **(D)** Representative gating strategy used to quantify the cell death status of mouse bone marrow 34^-^KSL cells at day +7 after lonafarnib treatment using 7-AAD/Annexin-V staining and quantification. Representative gating strategy used to quantify the cell death status of human **(E)** bone marrow or **(F)** cord blood CD34+/CD38-HSPCs at day +7 after lonafarnib treatment using 7-AAD/Annexin-V staining and quantification. **(G)** Flow cytometric quantification of p53 phosphorylation in cells at day +7 after culturing 34-KSL cells with lonafarnib; mean fluorescence intensity was normalized to vehicle levels. (*n=2-3 replicates from n=2 unique human donors/cell source from n=3 independent experiments for human studies; error bars show SEM, statistics show two-way ANOVA followed by Holm-Sidak’s multiple comparisons tests against vehicle*).

Previous work showed that inhibition of farnesyltransferase activity can cause cell death [13], leading us to consider that depletion of HSPCs after lonafarnib treatment may occur via changes in cell death status. Upon lonafarnib treatment, mouse 34^-^KSL cells were significantly more apoptotic and less live than vehicle treatment (**Figure 3D**). Human BM or CB CD34^+^ cells exhibited significantly more necrotic and apoptotic cells upon lonafarnib treatment compared to vehicle treatment (**Figure 3E-F**). Together, these data suggest that inhibiting farnesyltransferase activity causes HSPC cell death, consistent with prior findings showing that inhibition of farnesylation promote red blood cell death in zebrafish [13]. To better understand how this occurs, we quantified p53 activation in mouse 34^-^KSL cells after lonafarnib treatment. We detected a significant elevation in p53 activation upon lonafarnib treatment compared to vehicle-treated cells (**Figure 3G**). These data suggest that farnesyltransferase activity is critical for supporting the survival of HSPCs and implicate a potential role for p53 signaling in this process.

While certain lipids, like cholesterol, have been shown to affect hematopoiesis and hematopoietic cells [18, 19], the role of farnesylation in hematopoiesis is poorly understood. Our work suggests that farnesyltransferse activity is critical for the *ex vivo* expansion of human and mouse HSCs. Future work focused on defining the precise mechanisms by which this occurs and whether these pathways can be leveraged to better support HSC expansion are needed to better integrate these macromolecules into current paradigms of hematopoiesis. Pharmacological inhibition of farnesyltransferase in murine BM HSCs reduced cell expansion and increased cell death in hematopoietic stem and progenitor cell populations compared to control conditions. Intracellular levels of p53, a key tumor suppressor that plays a crucial role in both cell cycle regulation and apoptosis was elevated in murine HSCs following treatment. Our data suggest that farnesyltransferase activity is critical for HSC survival, *ex vivo* expansion ability, and HSC pool maintenance. A deeper understanding of how the metabolism of isoprenoids regulates HSC homeostasis can aid in developing therapeutics to support or restore hematopoietic system function in cases of disease. Finally, future work studying how we can leverage the MVA pathway to better support the *ex vivo* culture of HSCs is needed for us to optimize lipid metabolism to expand the availability of functional HSCs for life-saving treatments to patients who depend on them.

## ACKNOWLEDGMENTS

We would like to acknowledge Nicollette Setiawan, Taylor Billings, and Alex Hastie for their technical and administrative support of the project.

