## Supplemental Material for "Inhibition of farnesyltransferase activity diminishes hematopoietic stem cell *ex vivo* expansion ability"

**SUPPLEMENTAL METHODS**

**Mice:** Animal procedures were performed in accordance with the Fred Hutchinson Cancer Center Institutional Animal Care & Use Committee (PROTO2100049). Mice were housed and maintained in the Fred Hutch Comparative Medicine facility. Mixed-sex C57BL/6J or SJL mice (The Jackson Laboratory) aged 8-12 weeks were used for all studies.

**Analysis of scRNAseq datasets**: Previously published datasets were analyzed for the expression of Fnta/FNTA or Fntb/FNTB (1, 2). Previously published scRNAseq datasets were obtained from the Gene Expression Omnibus (GEO) under accession codes GSE207412 (mouse) and GSE139369 (human). Raw count matrices as provided by the original authors were downloaded and analyzed separately using R (version 4.4.0) and the Seurat package (version 5.2.0). No additional quality control or filtering steps were applied beyond those described in the original publications. Standard Seurat workflows were followed with default parameters as described in the Seurat tutorial (https://satijalab.org/seurat/articles/pbmc3k_tutorial.html). Briefly, data were normalized using LogNormalize with a scale factor of 10,000, followed by identification of variable features, scaling, principal component analysis (PCA), graph-based clustering, and visualization with Uniform Manifold Approximation and Projection (UMAP). Overall clustering and cell-type distribution were displayed using DimPlot. Author-provided cell type annotations were used for cell population labels. Expression of Fnta/fnta and Fntb/fntb was visualized using the Seurat FeaturePlot function (3), with expression values restricted to the 10th–90th percentile range to reduce the influence of outliers and noise.

**Mouse HSC extraction and analyses:** Mouse bone marrow (BM) cells were isolated from femurs and tibia, lysed with ACK buffer, and processed in complete IMDM (Gibco IMDM + 10%FBS + 1% penicillin-streptomycin). For hematopoietic stem/progenitor cell analyses, BM was lineage-depleted (Direct Lineage Depletion Kit, Miltenyi Biotec). Cells were stained with lineage antibody cocktail-V450, Sca1-BV605, cKit-APC-H7, CD34-PE, and 7AAD and FACs-sorted (BD FACSymphony S6). 5x10^2^ c-Kit^+^Sca-1^+^Lineage^-^CD34^-^ cells (34^-^KSL) were cultured in complete IMDM supplemented with mouse TPO, SCF, and FLT3 (TSF media), as described (4). For farnesyltransferase inhibition studies, media was supplemented with 0.1 µM or 1µM lonafarnib (Cayman Chemical, #11746) or DMSO vehicle control. Cultured cells were stained, washed, and analyzed by flow cytometry (BD FACS LSRFortessa X-50 or BD FACSymphony A5). For intracellular flow analysis, cells were antibody stained for surface markers and then fixed and permeabilized (BD Kit, 554714) before staining with phospho-p53 (Cell Signaling, 9284) or farnesyl (Invitrogen, PA1-12554). Data were analyzed using FlowJo.

***In vitro* human CD34^+^ cell analyses:** Human CD34^+^ cells derived from cord blood or BM used in all functional assays were purchased from STEMCELL Technologies and thawed according to the manufacturers protocol. 5x10^2^ cells were cultured in human TSF media supplemented with 0.1µM or 1µM lonafarnib or DMSO vehicle control. Cells were stained with human HSPC antibodies and analyzed using flow cytometry.

**Methylcellulose colony-formation assays:** 250 CD34^-^KSL mouse cells or 1000 human CD34^+^ cells were plated in methylcellulose medium (STEMCELL Technologies) supplemented with lonafarnib or DMSO vehicle control. Colonies were imaged and scored using a STEMVision system.

**Statistics**: Statistics were performed using Prism. Hypothesis tests between multiple groups or timepoints were performed using one- or two-way ANOVA followed by Holm-Sidak-corrected t-tests; tests comparing only two groups were performed using unpaired t-tests. Unless otherwise noted, data are considered significant when p<0.05.

**Supplementary Table 1: Manuscript Resources table.** Vendor information for antibodies, assays, mouse models, and software used throughout the manuscript.

| **ANTIBODIES** | | | | | | |
| --- | --- | --- | --- | --- | --- | --- |
| **Name** | | **Vendor** | **Catalog Number** | | **Clone** | **Dilution** |
| V450 Mouse Lineage Antibody Cocktail with Isotype | | BD Biosciences | 561301 | | n/a | 20µL / 10^6^ cells |
| APC/Cyanine 7 anti-mouse Ly-6A/E | | Biolegend | 108125 | | D7 | 1µg / 10^6^ cells |
| APC/Cyanine 7 Rat IgG2a, κ Isotype control | | Biolegend | 400523 | | RTK2758 | 1µg / 10^6^ cells |
| PE anti-mouse CD117 | | Biolegend | 105807 | | 2B8 | 1µg / 10^6^ cells |
| PE Rat IgG2b, κ Isotype control | | Biolegend | 400607 | | RTK4530 | 1µg / 10^6^ cells |
| PE/Cyanine 7 anti-mouse CD34 | | Biolegend | 119325 | | MEC14.7 | 1µg / 10^6^ cells |
| PE/Cyanine 7 Rat IgG2a, κ Isotype control | | Biolegend | 400521 | | RTK2758 | 1µg / 10^6^ cells |
| Purified Rat anti-mouse CD16/CD32 (Mouse BD FC Block) | | BD Biosciences | 553142 | | 2.4G2 | 2.5µg / 10^6^ cells |
| FITC Annexin V | | BD Biosciences | 556419 | | n/a | 5µl/test |
| FITC Arm Hamster IgG/Rat IgGb/RatIgG2a | | Biolegend | 78023 | | n/a | 5µl/test |
| Pacific Blue anti-human CD38 | | Biolegend | 356628 | | HB-7 | 2µl/test |
| Pacific Blue Mouse IgG1, k Isotype control | | Biolegend | 400151 | | MOPC-21 | 5µl/test |
| PE anti-human CD34 | | Biolegend | 343606 | | 561 | 2µl/test |
| PE Human IgG1 Isotype Control | | Biolegend | 403503 | | QA16A12 | 5µl/test |
| APC-H7 Mouse anti-human CD45RA | | BD Biosciences | 561212 | | 5H9 | 2µl/test |
| APC/Cyanine7 Mouse IgG2b, k Isotype Control | | Biolegend | 400328 | | MPC-11 | 5µl/test |
| APC anti-human CD90 (Thy1) | | Biolegend | 328114 | | 5E10 | 2µl/test |
| APC Mouse IgG1, k Isotype Control | | Biolegend | 400122 | | MOPC-21 | 5µl/test |
| Farnesyl Polyclonal Antibody | | Thermo Fisher Scientific | PA1-12554 | | n/a | 2µl/test |
| Phospho-p53 (Ser15) | | Cell Signaling Technology | 9284 | | n/a | 5µl/test |
| **Primary cell sources** | | | | | | |
| **Name** | **Vendor** | | | **Catalog Number** | | |
| Human BM CD34+ cells | STEMCELL Technologies | | | 70002.2 | | |
| Human CB CD34+ cells | STEMCELL Technologies | | | 70008.1 | | |
| **MOUSE MODELS** | | | | | | |
| **Name** | | **Vendor** | | | **Catalog Number** | |
| C57BL/6J | | The Jackson Laboratory | | | Strain #:000664 | |
| B6.SJL-Ptprca Pepcb/BoyJ | | The Jackson Laboratory | | | Strain#: 002014 | |
| **REAGENTS** | | | | | | |
| Iscove's Modified Dulbecco's Medium | | Gibco | | | 12440-053 | |
| Penicillin-Streptomycin Solution | | Fisher Scientific | | | MT30002CI | |
| Fetal bovine serum | | Fisher Scientific | | | 501527078 | |
| Compensation beads | | Fisher Scientific | | | 3031-5813-28 | |
| 7AAD cell viability stain | | BD Biosciences | | | 559925 | |
| Methocult GF M3434 | | STEMCELL Technologies | | | 3434 | |
| SmartDish | | STEMCELL Technologies | | | 27371 | |
| Ammonium-Chloride-Potassium (ACK) Lysing Buffer | | Fisher Scientific | | | 50-983-219 | |
| Mouse recombinant TPO | | STEMCELL Technologies | | | 78072 | |
| Mouse recombinant Flt/Flk-2 Ligand | | STEMCELL Technologies | | | 78011 | |
| Mouse recombinant SCF | | STEMCELL Technologies | | | 78064 | |
| Human recombinant Flt3/Flk-2 Ligand | | STEMCELL Technologies | | | 78009.1 | |
| Human recombinant TPO | | STEMCELL Technologies | | | 78210 | |
| Human recombinant SCF | | STEMCELL Technologies | | | 78062 | |
| Lonafarnib | | Caymen Chemical | | | 11746 | |
| **COMMERCIAL KITS** | | | | | | |
| **Name** | | **Vendor** | | | **Catalog Number** | |
| Direct lineage cell depletion kit mouse | | Miltenyi | | | 130-110-470 | |
| LS columns | | Miltenyi | | | 130-042-401 | |
| RNeasy Micro Kit | | Qiagen | | | 74004 | |
| **SOFTWARE** | | | | | | |
| **Name** | | **Vendor** | | | **Version** | |
| FlowJo | | BD Biosciences | | | 10 | |
| GraphPad Prism Version | | GraphPad Software | | | 10.4.1 (532) | |
| STEMVision Analyzer | | STEMCELL Technologies | | | 2.6.7.0 | |
| STEMVision Colony Marker | | STEMCELL Technologies | | | 2.5.0.0 | |
| R | | R foundation for statistical computing | | | 4.4.2 | |
| Seurat | | Hao et al. Nature Biotech 2023 | | | 5.2.0 | |
